# Diversity, function and evolution of marine invertebrate genomes

**DOI:** 10.1101/2021.10.31.465852

**Authors:** Yaolei Zhang, Jiahao Wang, Meiqi Lv, Haoyang Gao, LingFeng Meng, A Yunga, Inge Seim, He Zhang, Shanshan Liu, Linlin Zhang, Xin Liu, Xun Xu, Huanming Yang, M10K+ Consortium, Guangyi Fan

**Affiliations:** BGI-Qingdao, BGI-Shenzhen, Qingdao 266555, China; BGI-Shenzhen, Shenzhen 518083, China; China National GeneBank, BGI-Shenzhen, Shenzhen, 518120, China; Integrative Biology Laboratory, College of Life Sciences, Nanjing Normal University, Wenyuan Road 1, Nanjing 210046, China; Institute of Oceanology, Chinese Academy of Sciences, Qingdao 266071, China; James D. Watson Institute of Genome Sciences, Hangzhou 310058, China

## Abstract

Invertebrates, animals (metazoans) without backbones, encompass ∼97% of all animal yet remains understudied. They have provided insights into molecular mechanisms underlying fundamentally identical mechanisms in phylogenetically diverse animals, including vertebrates. Marine invertebrates have long fascinated researchers due to their abundance, diversity, adaptations, and impact on ecosystems and human economies. Here, we report a compendium and appraisal of 190 marine invertebrate genomes spanning 21 phyla, 43 classes, 92 orders, and 134 families. We identify a high proportion and long unit size of tandem repeats, likely contributing to reported difficulties in invertebrate genome assembly. A well-supported phylogenetic tree of marine invertebrates from 974 single-copy orthologous genes resolved topological controversies. We show that Ctenophora is at the basal phylum and Porifera is the sister group of Parahoxozoa; that Xenacoelomorpha is within Bilateria and is the sister group to Protostomia, rejecting three out of four hypotheses in the field; and that Bryozoa is at the basal position of Lophotrochozoa, not grouped into Lophophorata. We also present insights into the genetic underpinnings of metazoans from *Hox* genes, innate immune gene families, and nervous system gene families. Our marine invertebrate genome compendium provides a unified foundation for studies on their evolution and effects on ecological systems and human life.

## Introduction

Invertebrates, encompassing ∼97% of all animals (*1*), are named such because they lack a vertebral column (i.e., ‘backbone’). With high diversity in morphology, invertebrates consist of 32 phyla, of which marine invertebrates occupy 31. Marine invertebrates include a total of 178,023 extant species (*2*) (Data from The World Register of Marine Species (WORMS, 1-7-2020)) (**Figure S1**). These include ctenophores, sponges, corals, jellyfishes, sea stars, sea slugs, shrimps, crabs, squids, and oysters – many of which have crucial roles in ecosystems. Nine major phyla occupy 97.17% of all marine invertebrate species: Arthropoda, Mollusca, Annelida, Platyhelminthes, Cnidaria, Porifera, Echinodermata, Bryozoa, and Nematoda. Given their abundance, diversity, and impact on ecosystems and human economies, marine invertebrates have long fascinated researchers. Some marine invertebrates are harmful to humans, economically important animals, and plants. Others provide benefits. For example, marine species are harvested or cultured for human consumption, and some molluscan species can supply nacre (‘mother-of-pearl’). Despite their importance, invertebrate species have gathered relatively little attention compared to vertebrates and plants (*3*) and have by many been considered ‘lower’ or more ‘primitive’ lifeforms (*4*).

In the past two decades, increasingly powerful genome sequencing technologies and analyses have fuelled significant advances in our understanding of economically important species [i.e., fish, livestock, and plants] and allowed unprecedented insights into the evolution of life. In contrast, invertebrate genomics research has primarily employed a more ‘traditional’ lens, focusing on morphology limited molecules or genes. Efforts such as the Global Invertebrate Genomics Alliance (GIGA)(*5*) and BGI-Qingdao’s International Conference on Genomics of the Ocean have begun to fill this research gap. An improved understanding of marine invertebrate genomes is sure to enhance our ability to decrease their negative impacts and provide positive benefits to ecological systems and human life. Research on invertebrates also provides insights into the immune system, the nervous system, and body plan of animals at the most fundamental level. To further this effort, we present an appraisal of marine invertebrate genomes, highlighting salient themes from the unified data.

## Results

### Marine invertebrate genomes

Although the number of marine invertebrate species is enormous, only 190 species have a whole-genome assembly to date (data as of 1-7-2020) – far fewer than the number of sequenced vertebrates (approximately 700 species). The 190 species cover 21 phyla, 43 classes, 92 orders, and 134 families. Arthropoda (46 species) and Mollusca (51 species) represent more than a half, followed by Cnidaria (35 species), Echinodermata (19 species), and Rotifera (10 species) (**Figure 1 and Table S1**). The first sequenced marine invertebrate was the purple sea urchin (*Strongylocentrotus purpuratus*) sequenced in 2005 (*6*). Unfortunately, the following decade was marked by little progress in marine invertebrate sequencing. The development of next-generation sequencing technologies from the mid-2010s and a dramatic reduction in sequencing cost provided new marine invertebrate research opportunities. The third wave of sequencing technologies, since 2017, fuelled the generation of a large number of invertebrate genome assemblies. The number of sequenced invertebrate species now accounts for more than half of all sequenced species (**Figure S2**), a remarkable turnaround in the past two years.

**Figure 1.**
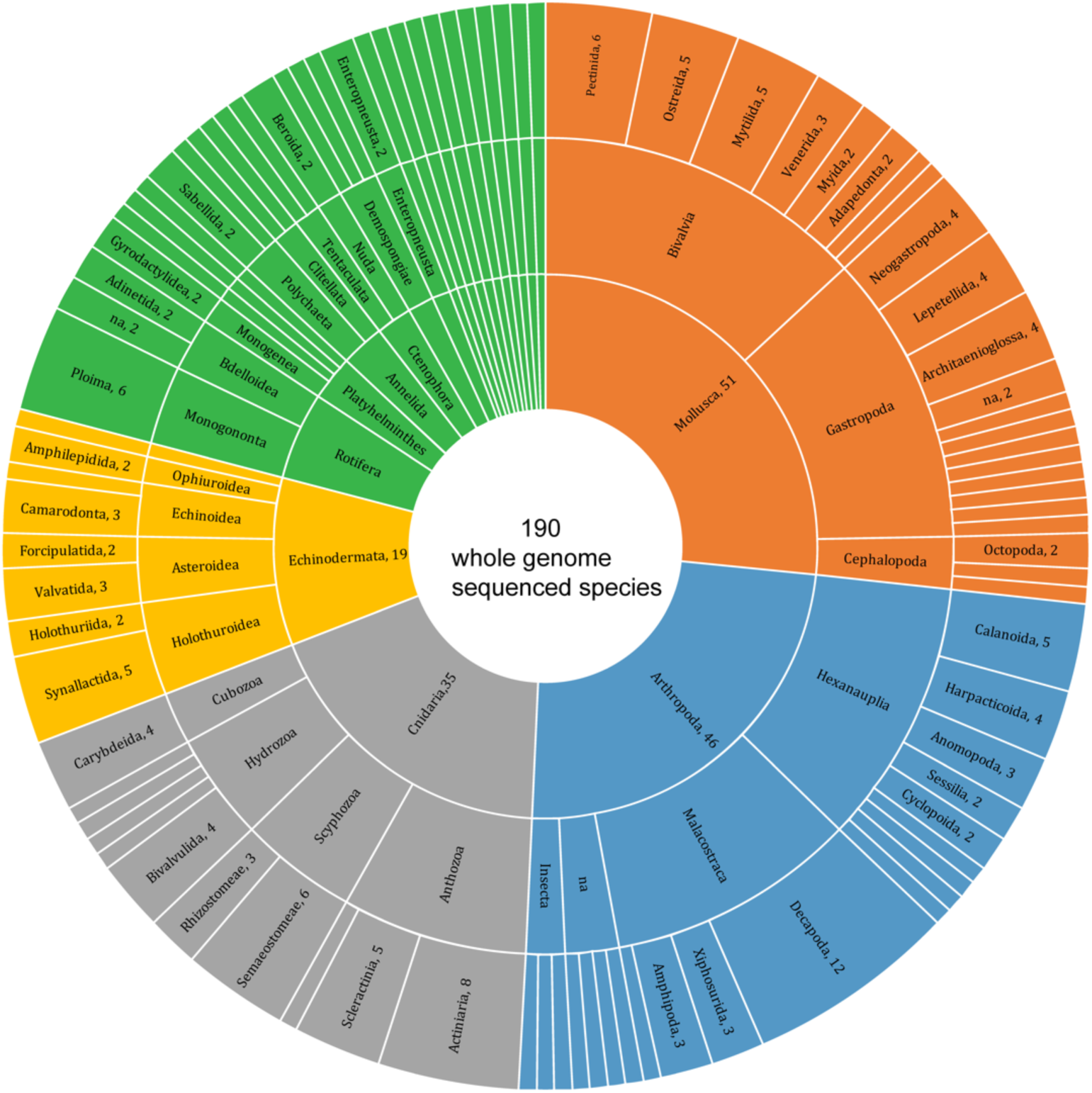
Distribution of 190 marine invertebrate species with nuclear genome assemblies.

The high diversity of marine invertebrate manifests as a broad spectrum of genome sizes. About ∼91% of sequenced marine invertebrate species have genome sizes smaller than 2 Gb (**Figure 2, Figure S3** and **Table S1**). The genome size of 62 species is smaller than 300 Mb (the smallest vertebrate genome, that of the fugu, is ∼360 Mb (*7*) and considered an extreme outlier), and 18 genomes are smaller than 100 Mb (**Figure 2, Figure S3** and **Table S1**). Four species have compact, ∼3-4 Mb genomes: the sponge *Aplysina aerophoba*, the shipworm *Bankia setacea*, and the copepods *Calanus glacialis* and *C. finmarchicus*. These species are found in three phyla, suggesting that they independently evolved remarkably compact genomes. There are only six species (five Arthropoda and one Mollusca species) with larger than 3 Gb genomes. The five Arthropoda species are found in the crustacean order Decapoda. The genomes of the shrimp *Palaemon carinicauda* (Arthropoda) and the squid *Euprymna scolopes* (Mollusca) are about 6.70 and 4.81 Gb, respectively. In contrast to many vertebrate genomes, where there is a strong correlation between genome size and repeat content (see detail in “Review of Marine Tetrapods” paper preprinted at the same time), invertebrate genomes do not show an apparent linear relationship between genome size and repeat content (**Figure 2**). For example, the parasitic worm *Pomphorhynchus laevis*, one of ten species with a genome repeat content larger than 60%, is ∼260 Mb genome and consists of 64.0% repeats. At the other extreme, the crayfish *Procambarus virginalis* has a 3.3 Gb genome, but only 8.8% is repeats (**Figure 2**). Of course, some species have both large genome sizes and high percentages of repeats. For example, the blue king crab (*Paralithodes platypus*) has a 4.8 Gb genome with 77.2% of repeats. We also found that seven species have less than 10% repeats in their genomes. The proportion of repeat subtypes is also diverse (**Figure S4**). Approximately ∼90.53% of species have a low proportion of SINEs (less than ∼3%). Unclassified repeats constitute more than half of the repeat content in five phyla (Rotifera, Arthropoda, Echinodermata, Platyhelminthes, and Porifera). Marine invertebrate genomes also have quite diverse GC contents. The average GC content is about 36.31%, which is slightly lower than that of marine mammals (38.2%) and higher than that of fish (32.8%). Outliers include six sequenced species with GC content higher than 47% – 59.6% in the case of the sponge (*Aplysina aerophoba*) – and 26 species with GC content less than 30% (**Figure S3**).

**Figure 2.**
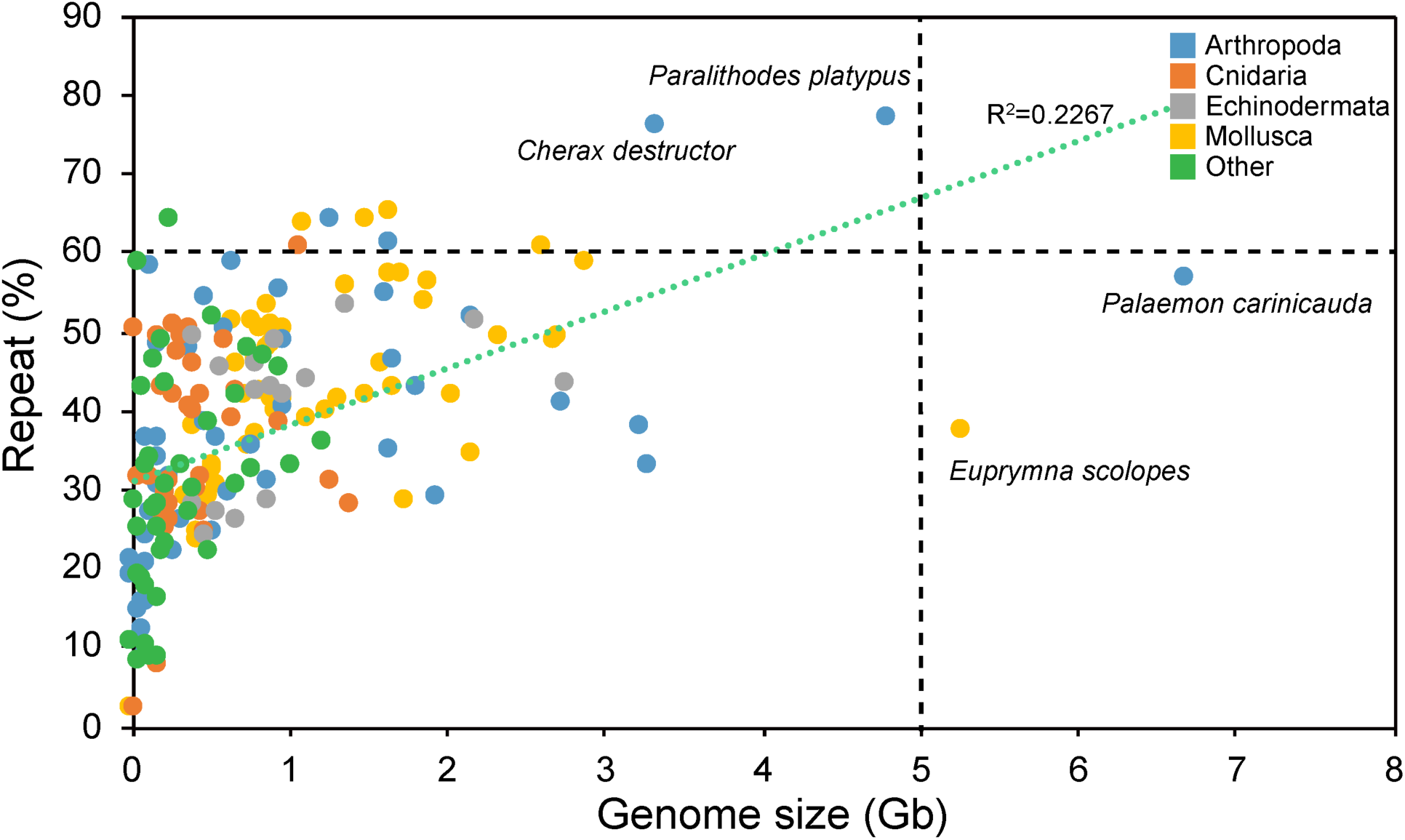
Genome sizes and repeat contents of whole-genome sequenced marine invertebrates (190 species). There is no obvious correlation between genome size and repeat content (*R*^*2*^=0.2267).

It becomes apparent that the various genomic features of marine invertebrates can contribute to genome assembly difficulty. Only 22 genomes out of 190 have been assembled at the chromosome level. Of these, about a quarter (47 genomes) have a scaffold N50 less than 20 kb – far below current gold standards for genome annotation. Moreover, of the 143 genomes with a scaffold N50 larger than 20 kb, some have a low contig N50 –18 less than 10 kb, five as low as 3.7 kb – indicating that many assemblies have numerous gaps and likely incomplete gene models.

### Marine invertebrate phylogeny

To resolve marine invertebrate phylogeny, we constructed trees based on a single-copy orthologous gene set from the 190 whole-genomes assemblies covering 21 phyla. Genomes with scaffold N50 larger than 20 kb were deemed adequate for identifying single-copy orthologous groups (SCOGs), retaining 143 genomes from 20 phyla. To include the phylum Nematoda in our phylogenetic analysis, we included the most complete marine genome from this phylum assembled to date: the parasitic herring worm (*Anisakis simplex*) (scaffold N50 ∼6.8 kb, NCBI accession: GCA_900576815.1). As discussed for a long time, tree reconstruction errors may be caused by inadequate models,long branch attraction, short internal branches, and especially unrepresentative data. Therefore, here we identified SCOGs by interrogating genome assemblies using BUSCO (v2.0) (Benchmarking Universal Single-Copy Orthologue) with the metazoan gene database (*8*). A total of 974 SCOGs (BUSCO genes) were identified in the 190 genomes. We excluded four species with low BUSCO single-copy ortholog completeness scores: *Bankia setacea* (Mollusca; 3.7%), *Anentome helena* (Mollusca; 11.5%), *Aplysina aerophoba* (Porifera; 3.5%), and *Kudoa iwatai* (Cnidaria; 9.5%). The final orthologous gene set included 140 species in 21 phyla (average BUSCO completeness score 70.7%).

We used MAFFT (*9*) to perform protein sequence alignments for each SCOG and trimmed the results using Aliscore (*10*) and Alicut (*11*) (to remove ambiguously aligned regions with little to no phylogenetic signal), yielding a 365,588-amino acid alignment matrix. IQ-TREE (*12*), which can quickly test substitution models and choose the best model to generate reasonable trees, was employed to construct 974 gene trees. ASTRAL (*13*) was used to infer a consensus species tree (**Figure 3**). To further validate the constructed trees, we also used the popular tool RAxML (*14*) to build gene trees with the ‘PROTGAMMAILG’ model and 100 rounds of bootstrapping, followed by species tree inference using ASTRAL (**Figure 4**). Except for the position of Porifera, the RAxML and IQ-TREE trees were identical. To evaluate the robustness of our reconstructed phylogeny, we also considered the phylogenetic position of Placozoa, Xenacoelomorpha and Bryozoa with a subset of species. The species trees agreed (**Figure 5a, b, c, d, e** and **f**) with the topology estimated using 140 species. Three major findings can be inferred from the phylogeny. Firstly, Placozoa and Cnidaria are sister groups, resolving a longstanding controversy (*15-21*). Secondly, Xenacoelomorpha is nested within Bilateria and is the sister group to Protostomia, rejecting three out of four hypotheses in the field (*22-29*). Thirdly, Bryozoa is at the basal position of Lophotrochozoa, not grouped into Lophophorata. The phylogenetic position of Xenacoelomorpha (placed as a sister group to Nephrozoa) (**Figure 5g**), nor the position of Bryozoa within the ‘traditional’ Lophophorata (a grouping of bryozoans, brachiopods, and phoronids) (**Figure 5h** and **i**), agrees with a recent study by Cannon and colleagues (*29*) based on transcriptome data from 77 metazoan taxa [60 invertebrates, of which 55 are marine species and 18 overlapped our data set]. This controversy is possibly attributed to an erroneous assignment of paralogous genes as orthologs in the previous study and a limited overlap with our 974-gene BUSCO-derived set (115/974, 11.7%; and 309/974, 31.6%). Taken together, we propose that our inferred phylogeny **(Figure 5j)** is robust and a valuable resource for future studies.

**Figure 3.**
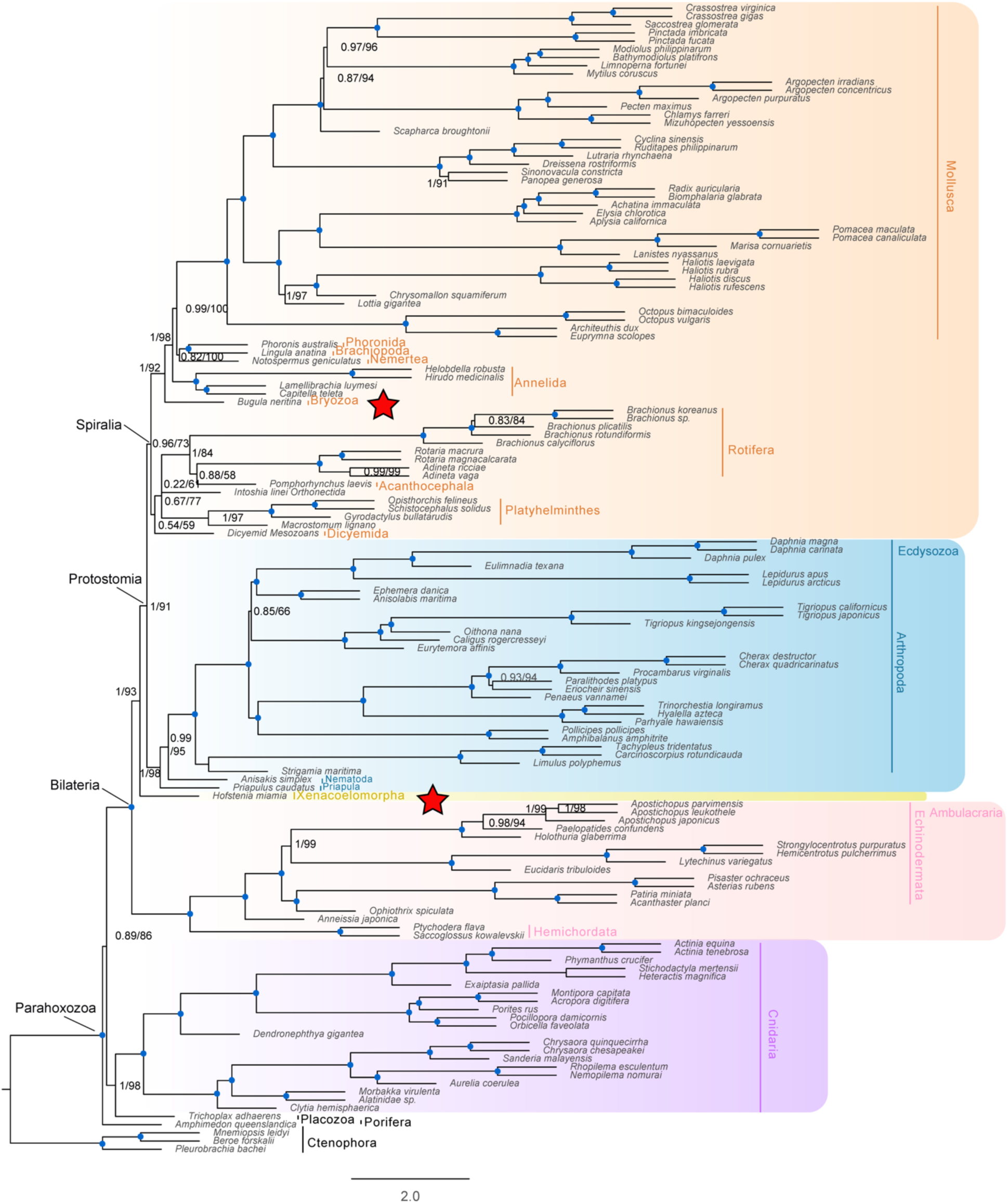
The maximum likelihood phylogenetic tree constructed using whole-genome sequences of 140 marine invertebrate species and IQ-TREE. The species tree was inferred using ASTRAL and 974-gene trees (IQ-TREE) of metazoan BUSCO genes. The numbers presented “*/*” near the branches represent reliability evaluation inferred by IQ-TREE (1 means 100% support) / bootstrap values. The blue dots represent these branched are “1/100” support. The red stars represent the different positions of important phyla inferred from this study compared to previous researches.

**Figure 4.**
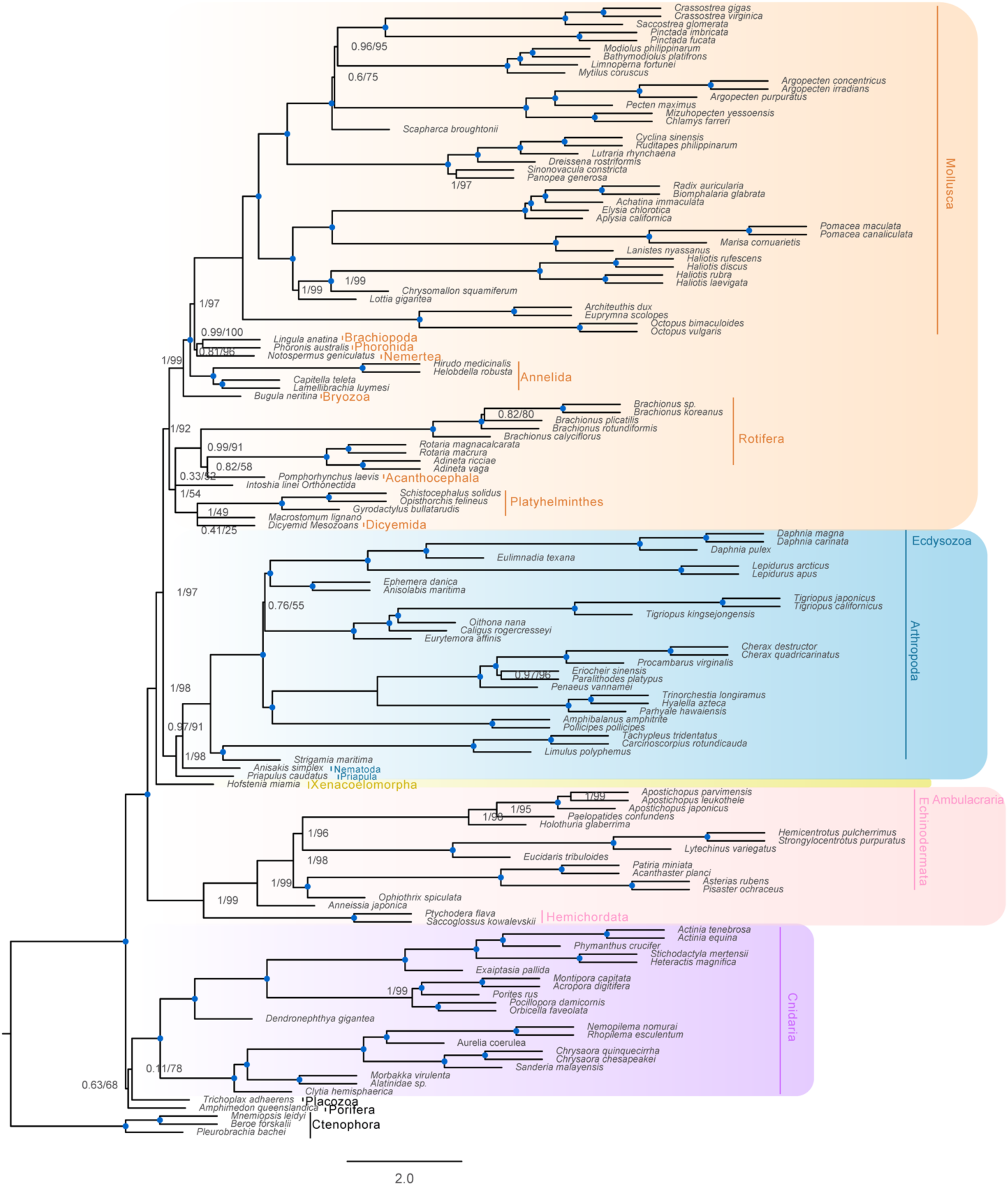
The maximum likelihood phylogenetic tree constructed using whole-genome sequences of 140 marine invertebrate species and RAxML. The species tree was inferred by using ASTRAL based on 974-gene trees (RAxML) of metazoan BUSCO genes.

**Figure 5.**
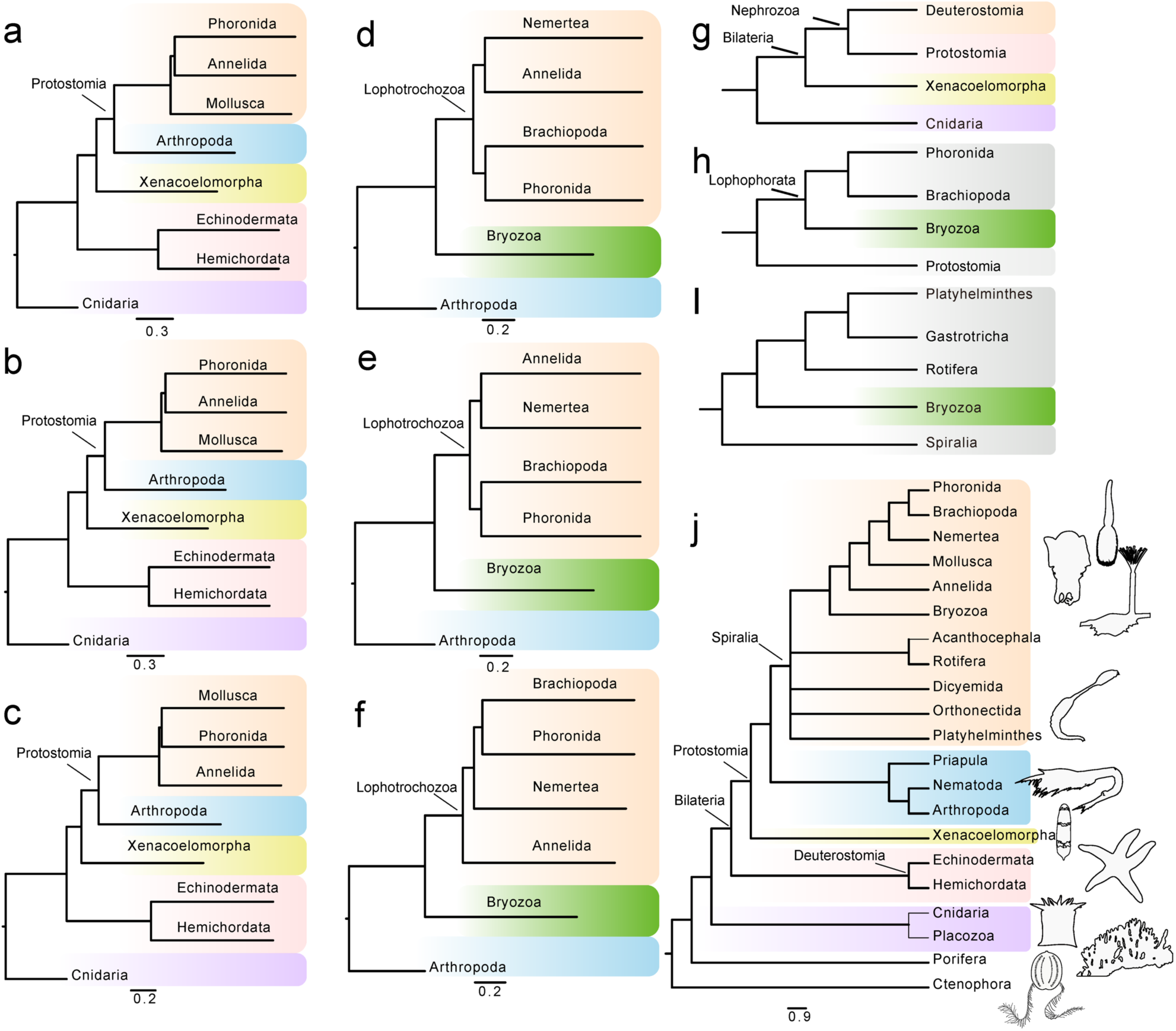
Verification of our constructed trees using subset species with different methods and inferred phylogenetic tree of 21 phyla by combining our data. a, b and c), phylogenetic trees for verification of Xenacoelomorpha position using IQ-TREE, RAxML and MrBayes, respectively. The three trees were constructed using eight representative species, including *Hofstenia miamia* (Xenacoelomorpha), *Pocillopora damicornis* (Cnidaria,), *Acanthaster planci* (Echinodermata), *Saccoglossus kowalevskii* (Hemichordata), *Daphnia magna* (Arthropoda), *Capitella teleta* (Annelida), *Pomacea maculate* (Mollusca) and *Phoronis australis* (Phoronida). d, e and f: phylogenetic trees for verification of Bryozoa position using IQ-TREE, RAxML and MrBayes (*49*), respectively. The three trees were constructed using six representative species, including *Bugula neritina* (Bryozoa), *Tigriopus californicus* (Arthropoda), *Capitella teleta* (Annelida), *Notospermus geniculatus* (Nemertea), *Lingula anatina* (Brachiopoda) and Ph*oronis australis* (Phoronida). g), Phylogenetic position inferred by Cannon et al (*29*). h), Bryozoa position inferred by Cannon et al. i) Bryozoa position from traditional view(*52*). j) inferred relationship of 21 marine phyla by combing all data in this study.

### Homeobox genes

Evolutionary biologists have long classified invertebrates into radially symmetrical and bilaterally symmetrical animals based on morphology. Radially symmetrical animals include the phyla Porifera (sponges), Cnidaria (jellyfishes, corals, and related species), Ctenophora (comb jellies), and Placozoa (show the simplest morphology and is one of the most basal metazoan phyla). The remaining 17 marine phyla analysed in this study are bilaterally symmetrical (i.e., possesses two orthogonal body plans) and are members of the evolutionary lineage Bilateria. Some species not grouped within Bilateria have bilateral symmetry (*30*), however. A salient example is *Nematostella vectensis* (starlet sea anemone) of phylum Cnidaria (*31*).

It now appreciated that the loss or expansion of genes encoding homeobox proteins (particularly the *Hox* genes), transcription factors that determine the identity of body segments, is key to the evolution of a bilaterian body. We used our 140-species whole-genome data set to investigate the homeobox gene repertoire of marine invertebrates (**Figure 6**). We identified very few homeobox genes or gene fragments in the basal phyla Ctenophora (*Hox2, Hox4, Hox5*, and posterior *Hox* genes), Placozoa (*Hox1, Hox6, Hox8*, and posterior *Hox* genes), and Porifera (*Hox2* and *Hox4*, and a *Hox7* fragment). Cnidaria genomes contain nine intact (*Hox2, Hox3, Hox4, Hox5, Hox6, Hox7, Hox8*, and posterior *Hox* genes) and one fragmented (*Hox1*) *Hox* genes. In the Bilateria lineage, *Hox1* is found in most phyla (85 out of 116 species). A lack of *Hox1* in Dicyemida, Bryozoa, Rotifera, Orthonectida, and Platyhelminthes suggests that bilateral symmetry can be achieved without this anterior *Hox* gene. The phylum Arthropoda includes Chelicerata (sea spiders and horseshoe crabs). We found multiple copies of the anterior *Hox* genes (*Hox1* to *Hox4*) in marine Chelicerata genomes, likely resulting from whole-genome duplication (WGD) and in agreement with recent work on the mangrove horseshoe crab (*Carcinoscorpius rotundicauda*) genome (*32*) and sea spider transcriptomes(*33*). Interestingly, cephalopods (phylum Mollusca) – giant squid (*Architeuthis dux*), Hawaiian bobtail squid (*Euprymna scolopes*), California two-spot octopus (*Octopus bimaculoides*), and common octopus (*Octopus vulgaris*) – have lost several homeobox genes, in particular anterior *Hox* genes (*Hox1, Hox2, Hox3*, and *Hox4*), in agreement with previous analyses of cephalopod *Hox* genes(*34*). Consistent with recent studies (*35*), we found that most Bivalvia species (20 out of 22) have lost *Hox7* (also known as *Antp*). While except for Rotifera, most genomes contain more than one gene copy of the *Hox8*/*AbdA*/*Utx*/*Lox2*/*Lox4* group (denoted HOX8 in **Figure 6**).

**Figure 6.**
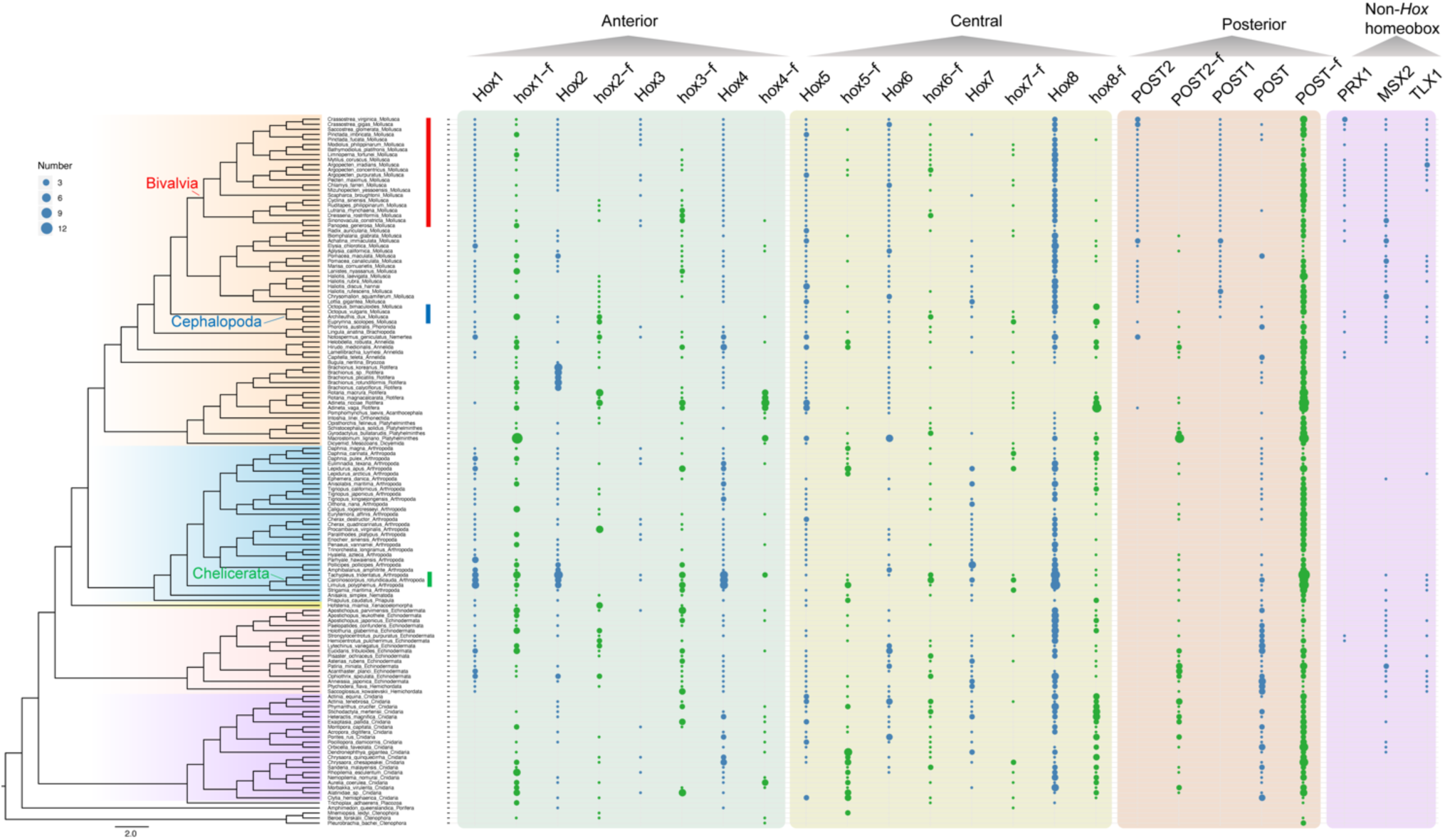
Overview of homeobox genes identified in 140 marine invertebrate species. The figure shows distribution of anterior (*Hox1* to *Hox4*), central (*Hox5* to *Hox8*) and posterior *Hox* genes (*POST, POST1* and *POST2*), as well as non-*Hox* homeobox genes (*PRX1, MSX2*, and *TLX1*). Blue dots indicate homeobox genes; green dots, homeobox fragments (less than 80 amino acids). The red, blue and green bars next to the species names indicate Bivalvia, Cephalopoda and Chelicerata species. Genes or gene fragments (with the suffix ‘-f) were assigned using a sigmoid model by calculating protein lengths of predicted gene (**Figure S5**).

### Innate immune gene families

Similar to plants, invertebrates lack an adaptive immunity but have developed a complex innate immune system(*36, 37*). We investigated the classical innate immune receptor repertoire of marine invertebrates (**Figure 7**). Six Toll-like receptors (TLRs) *TLR1, TLR2, TLR3, TLR4, TLR7*, and *TLR13* were identified. No TLRs were found in Ctenophora, Porifera, Placozoa, Cnidaria, Xenacoelomorpha, Platyhelminthes, Orthonectida, Acanthocephala, Rotifera, Dicyemida, Annelida, Nemertea, and the majority of Arthropoda. TRL genes were present in Bilateria, including Ambulacraria and Mollusca. In Ostreida and Pectinida species, five TLR genes were found. TLR1 was only found in Pectinida, indicative of an ancestral origin. *TLR3* (at least six copies) and *TLR13* (at least nine copies) were expanded in Pectinida. *TLR13* expanded in Ostreida [29 copies in Sydney rock oyster (*Saccostrea glomerata*), 31 copies in the eastern oyster (*Crassostrea virginica*) and 55 copies in the Pacific oyster (*Crassostrea gigas*)]. Four types of NOD-like receptors (NLRs) – including *NLRC3, NLRC5*, NLR Family Pyrin Domain Containing 12 (*NLRP12*), and *NLRP14* – were found in Porifera but lost in some species of Cnidaria, Echinodermata, Arthropoda, Platyhelminthes, Orthonectida and Mollusca. CARD Domain Containing 3 (NLRC3) genes were distributed among ten phyla and significantly expanded in Scleractinia species (25-135 copies). The scavenger receptors (SCARs) *SCARA1* (in particular), *SCARA2, SCARB1* and *SCARB2* are broadly distributed in marine invertebrates, especially in Cnidaria, Echinodermata and Mollusca. In contrast, SCARs could not be identified in Rotifera, Platyhelminthes and Arthropoda. Concurring with the literature(*38*), genes encoding peptidoglycan recognition proteins (PGRPs) are absent in radiation-symmetrical animals, including Ctenophora, Porifera, Placozoa and Cnidaria. Broadly, our results suggest that the innate immune system of radiation-symmetrical invertebrates employ NLRs and SCARs, hinting at a fundamental role of the immune system. In bilateral invertebrates, TLRs and PGRPs may have further enhance innate immunity. Arthropod species have lost most of the classic innate immune families examined here (TLRs, NLRs, SCARs, and PGRPs), suggesting that alternative components have evolved. Indeed, the expansion and robust expression of Dscam genes may endow the horseshoe crab with a strong immune and environmental adaptability (*39*). Rotifera species have lost SCARs but retained PGRPs and NLRC3, which may be associated with their freshwater habitats. A subset of Ambulacraria and Mollusca (Ostreida and Pectinida) genomes harbour all the four gene families examined and gene expansions (e.g., *TLR3, TLR4, TLR13, SCARA1*, and *SCARA2*), indicating that they have a robust immune system.

**Figure 7.**
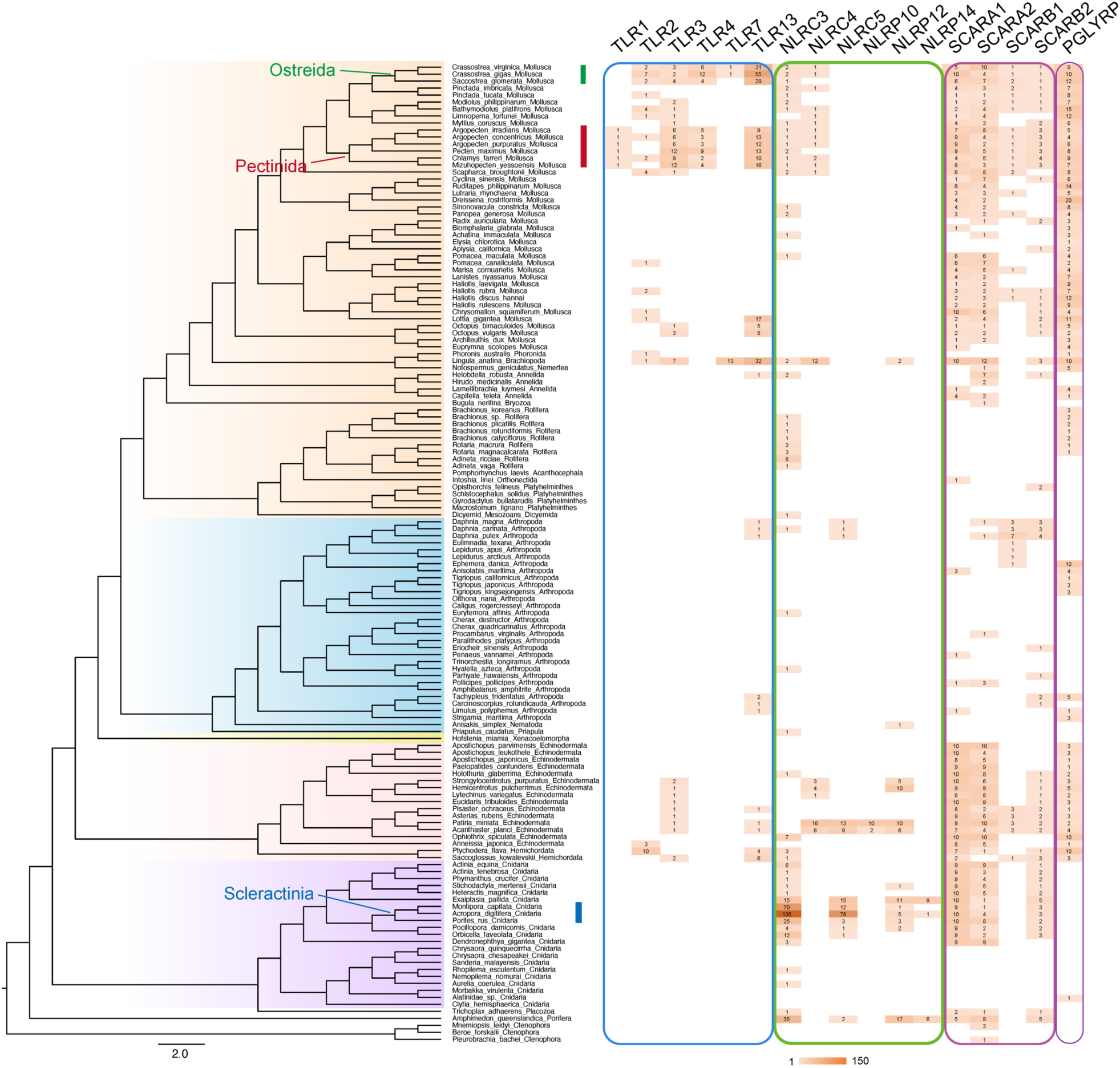
Distribution of four gene families of the innate system in marine invertebrates. The heat map shows the number of genes (the number is shown in the centre). The suffix TLR denotes Toll-like receptor genes; NRL, NOD-like receptor genes; SCAR, scavenger receptors genes; PGRPs, peptidoglycan recognition protein genes. The green, red and blue bars next to species names represent Ostreida, Pectinida and Scleractinia species.

### Nervous system gene families

Although the invertebrate nervous system is considered much simpler than those found in vertebrates, there is still a broad range in complexity. For examples, hydras and jellyfish have the simplest nervous system, termed a ‘nerve net’, without distinct central or peripheral regions. Sea stars have a further evolved nervous system, showing some centralised organisation. Some worms and molluscs evolved a distinct separation of the peripheral and central nervous systems. To investigate the potential evolutionary process of the marine invertebrate nervous system, we focused on three central neurodevelopment associated gene families: Neurogenins (NEUROGs), a family of bHLH transcription factors associated with neuronal differentiation; neurogenic differentiation factors (NEURODs), which are also involved in the differentiation of nervous system during early neural development; and protocadherins (PCDHs) which regulate neuronal development and mediate cell-cell adhesion. We identified three, three, and seven members of the NEUROG, NEUROD and PCDH families, respectively (**Figure 8** and **Fig. S6**). As expected, in the basal Ctenophora, Porifera and Placozoa, no or few genes were found.

**Figure 8.**
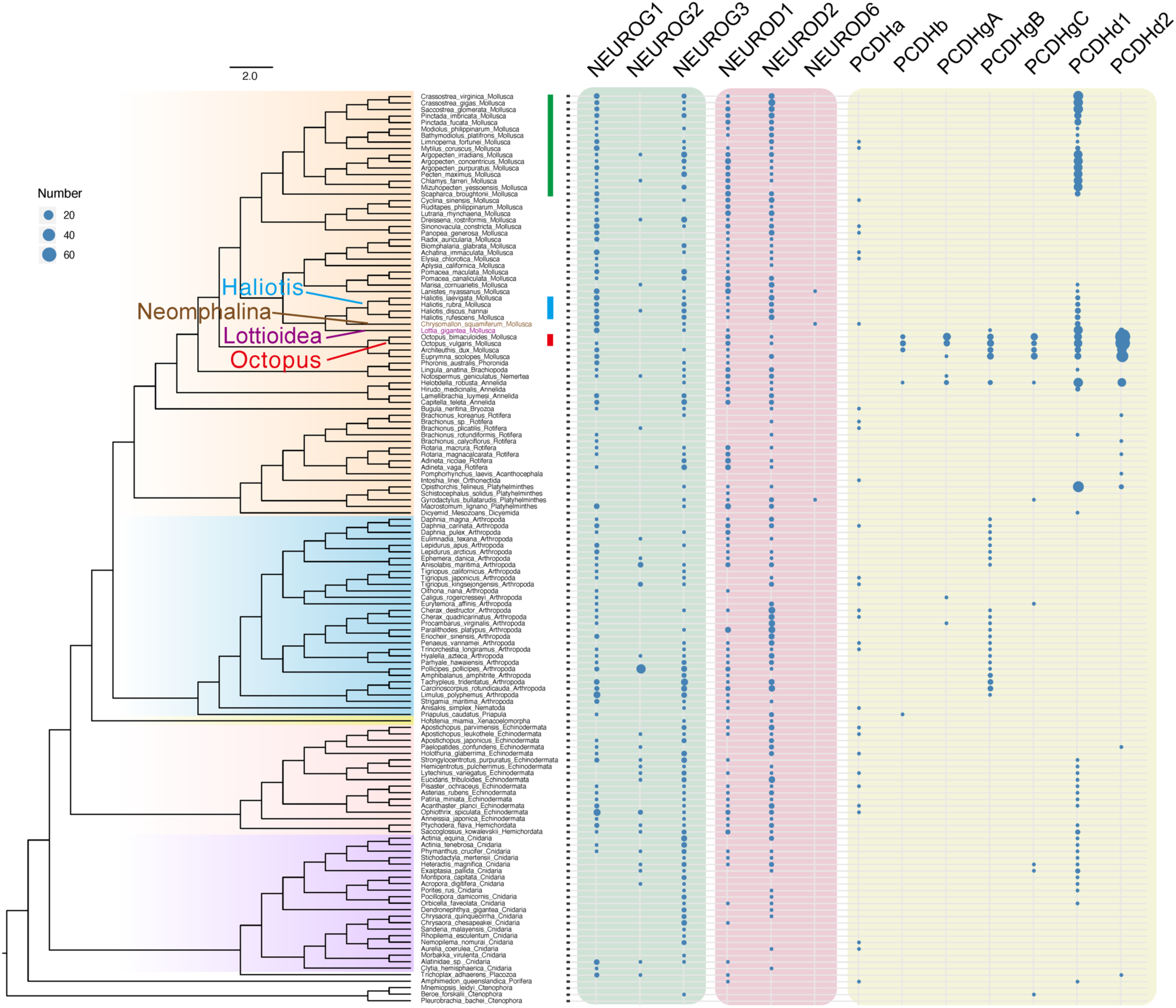
Distribution of gene families associated with nervous system development and function in marine invertebrates. The suffix NEUROG denotes Neurogenin genes; NEUROD, neurogenic differentiation factor genes; PCDH, protocadherin genes; The blue and red bars next to species names represent Haliotis and Octopus species.

In the NEUROG family, we found that *NEUROG1* is broadly distributed (81 species has *NEUROG1* and 11 species a *NEUROG1* fragment) – especially in Mollusca (37 out of 40 species), Arthropoda (24 out of 31 species) and Echinodermata (10 out of 15 species). For the remaining three, seven and five species for which *NEUROG1* was not identified, NEUROG2 or NEUROG3 were found in two, seven and five species, respectively, meaning that all Mollusca (except California two-spot octopus), Arthropoda and Echinodermata genomes contain at least one member of the NEUROG family. *NEUROG1* and *NEUROG2* were identified only in three and five Cnidaria species, while 19 out of 21 species were checked with *NEUROG3* (**Figure S6**). In the NEUROD family, the distribution of *NEUROD1* (identified in 38 out of 40 Mollusca species, 22 out of 31 Arthropoda species and 10 out of 15 Echinodermata species) was similar to *NEUROG1. NEUROG2* was identified in 30 species, while 74 harboured *NEUROG2* fragments. In contrast to neuroD/neurogenin, the distribution of PCDH gene family genes was phyla-specific. Protocadherin gamma subfamily B (P*CDHgB*) was mainly found in Arthropoda genomes, while protocadherin delta 1 (*PCDHd1*) was mainly found in part of Cnidaria (10 species), Ambulacraria (10 species), Platyhelminthes (1 species), Annelida (2 species) and Mollusca (27 species). Interestingly, all PCDH genes except PCDHa underwent a massive expansion in octopuses (phylum Mollusca). This includes 104 PCDHs in California two-spot octopus and 96 copies in common octopus, consistent with previous work (*34*). We identified 33 and 56 PCDHs in the giant squid and Hawaiian bobtail squid, respectively – agreeing with previous work showing a protocadherin gene expansion in this octopus sister group(*40, 41*).

## Discussion

More comprehensive research on marine invertebrates promises to answer a broad range of evolutionary, biological, and ecological questions. These include the origins of animal life such as the formation of multicellularity and early-animal evolution, nervous and immune system development, and ecological adaptations (e.g., biomineralisation and regeneration), breeding and aquaculture. High-throughput sequencing technologies provide us with an unprecedented opportunity to integrate traditional biological approaches with genomic data to describe new aspects of the evolutionary, functional and structural diversity of marine invertebrates. In this review, we constructed a database of marine invertebrate genomes, which currently includes 194 published genomes. This database will be continuously updated, including improving current assemblies using emerging sequencing technologies and assembly methods (e.g., see stLFR(*42, 43*)). While the current data set does not cover the full spectrum of marine invertebrate species (21 out of 31 phyla), and many of the genomes do not match current gold standards, we show that valuable insights can be gained.

The phylogenetic relationship of marine invertebrate has been contentious for a long time, especially in the case of Ctenophora, Porifera, Placozoa, Cnidaria and Xenacoelomorpha. Whole-genome assemblies enable the generation of orthologous gene sets that can be used to help resolve this dispute. Here, we used single-copy orthologous BUSCO genes to provide an unprecedented panorama of marine invertebrate phylogeny. We hope that this effort will be a valuable reference for future research. We also employed the data set to investigate the distribution of gene families associated with the immune system, the nervous system, and body plan of animals. Gene expansions generally agree with features observed in the immune and nervous system of invertebrates. NeuroD/neurogenin (*NEUROG1* and *NEUROD1*) appears to be correlated with a more complex nervous system in marine invertebrates, while an expansion of protocadherin (PCDH) genes likely underlies the exceptional intelligence of octopuses and squids. While our data support the concept of *Hox* genes underlying the invertebrate body plan and driving bilateral symmetry, it must nevertheless be appreciated that complex gene clusters may not be resolved in genome assemblies without chromosome-conformation-capture methods such as Hi-C(*44*).

In summary, the data and results presented in this review provides a valuable, continuously updated resource for and insights into marine invertebrates.

## Materials and Methods

### Genome data collection

Marine invertebrate species information (species number and classification) were collected from “The World Register of Marine Species (WORMS, 1-7-2020)” (*2*) and NCBI database. Assembled genomes were mainly collected from NCBI database (see Table S1).

### Genome repeat annotation

Repeat sequences of all collected genomes were identified using a unified pipeline. Firstly, transposable elements were searched using RepeatMasker (v4.0.5) (*45*) and RepeatProteinMask (v4.0.5) against the Repbase database(*46*) at the nuclear and protein levels, respectively. Secondly, RepeatModeler (v1.0.8) and LTR-FINDER (v1.0.6) (*47*) were used to performing *de novo* prediction and construct a custom transposable element database, which was used to predict transposable element using RepeatMasker again. Thirdly, tandem repeats were predicted by using Tandem Repeat Finder (v4.0.7) (*48*).

### Construction of phylogenetic trees

All of the trees constructed in the present study are based on protein sequences of Single Copy Orthologous Genes (SCOGs). SCOGs were identified using BUSCO (v2.0) (Benchmarking Universal Single-Copy Orthologue) with the metazoan gene database (*8*), searching the whole genome assemblies. Next, MAFFT (*9*) was used to carry out protein sequence alignments for each SCOG and Aliscore (*10*) and Alicut (*11*) were used to remove ambiguously aligned regions with little to no phylogenetic signal. Then IQ-TREE (*12*), RAxML (*14*) and MrBayes(*49*) were used to construct gene trees for each SCOGs. Finally, ASTRAL (*13*) was used to infer a consensus species tree.

### Identification of gene families

For Hox, representative immune and nervous systems related gene family identification, we downloaded all Hox genes from the NCBI database, TLRs, NLRs, SCARs, PGRPs, NEUROGs, NEURODs and PCDHs from the KEGG database, and carried out a manual check to remove poor sequences to generate reference protein library. Then we mapped the reference proteins to 140 collected genomes using BLAT (*50*) and predicted relevant genes using GeneWise (v2.4.1) (*51*) based on the BLAT results.

## Author contributions

G.F., X.L. and X.X. designed this project. Y.Z. performed data analysis including comparative genomics, phylogenetic relationships, and manuscript writing. J.W. participated in data collection, repeats annotation and figures preparation. M.L., H.G., L.M., and Y. A. carried out data collection and repeat annotation. I. S. helped to improve the manuscripts and supplied good suggestions. All authors read and approved the final manuscript.

## Supplementary Figures

**Figure S1.**
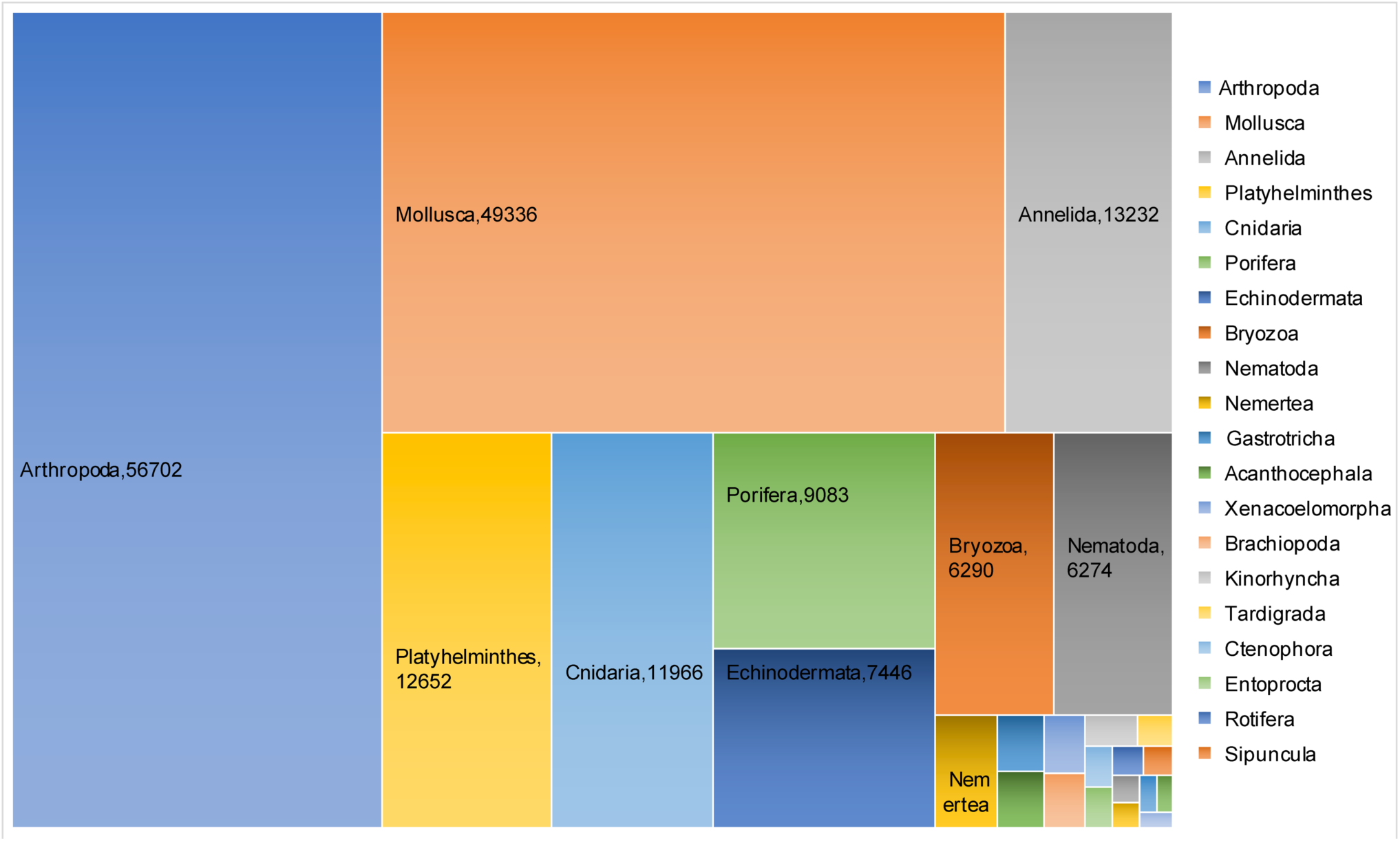
Classification and species numbers of marine invertebrates.

**Figure S2.**
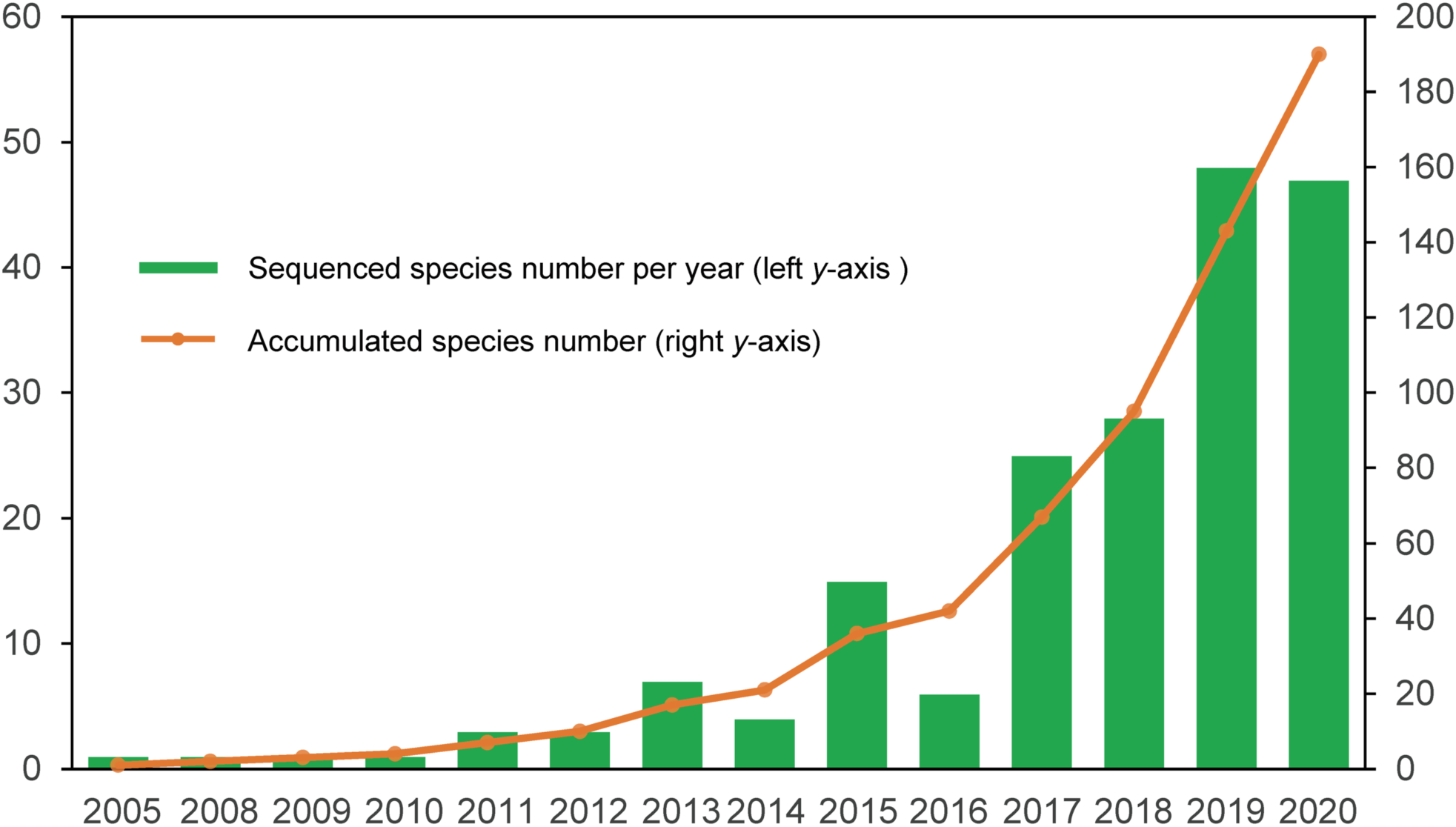
Summary of published marine invertebrate genomes since 2005.

**Figure S3.**
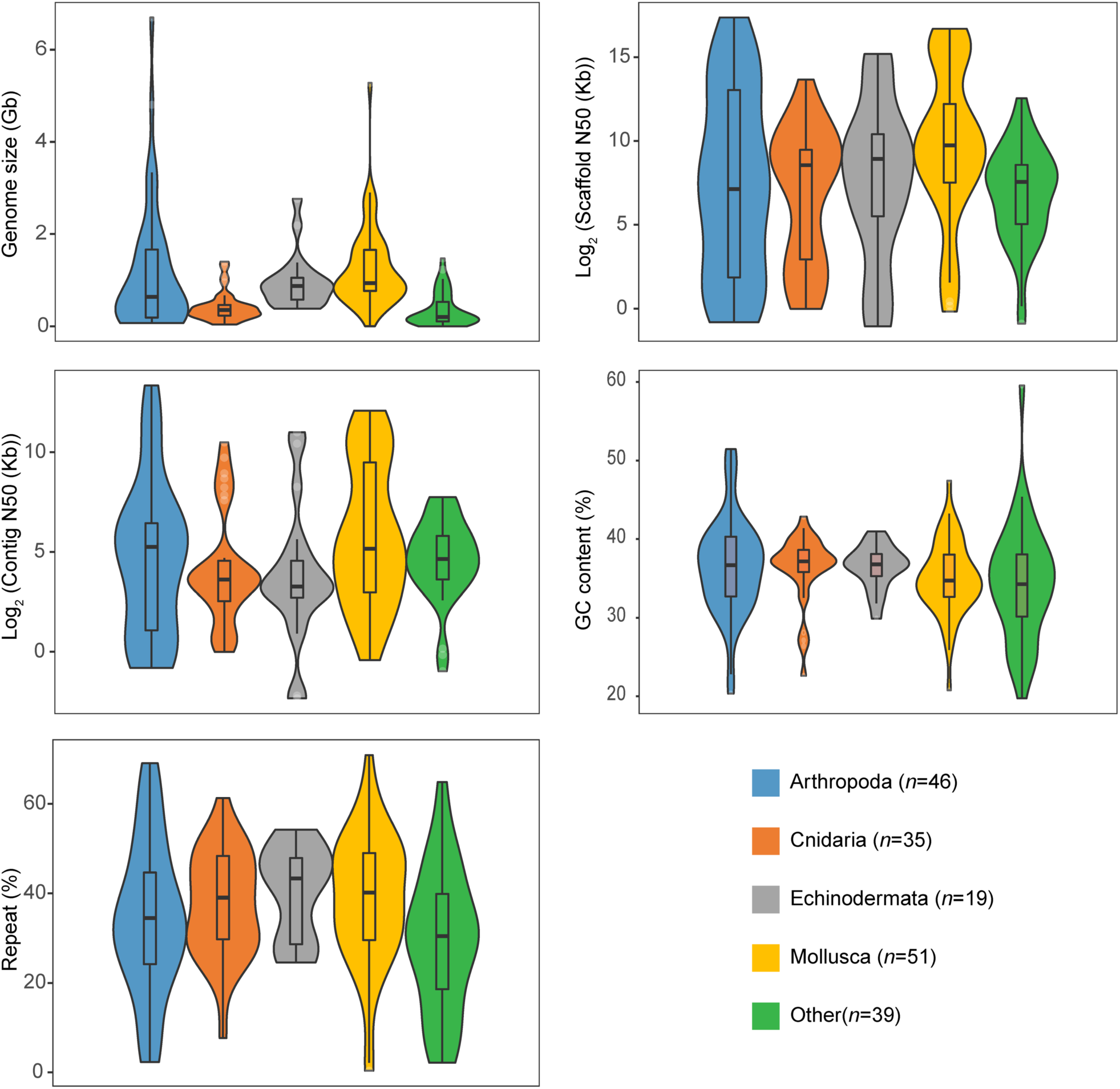
Summary of the genome features genome size, percent GC and repeats, and scaffold and contig N50 lengths (in kb) of marine invertebrates with a whole-genome assembly.

**Figure S4.**
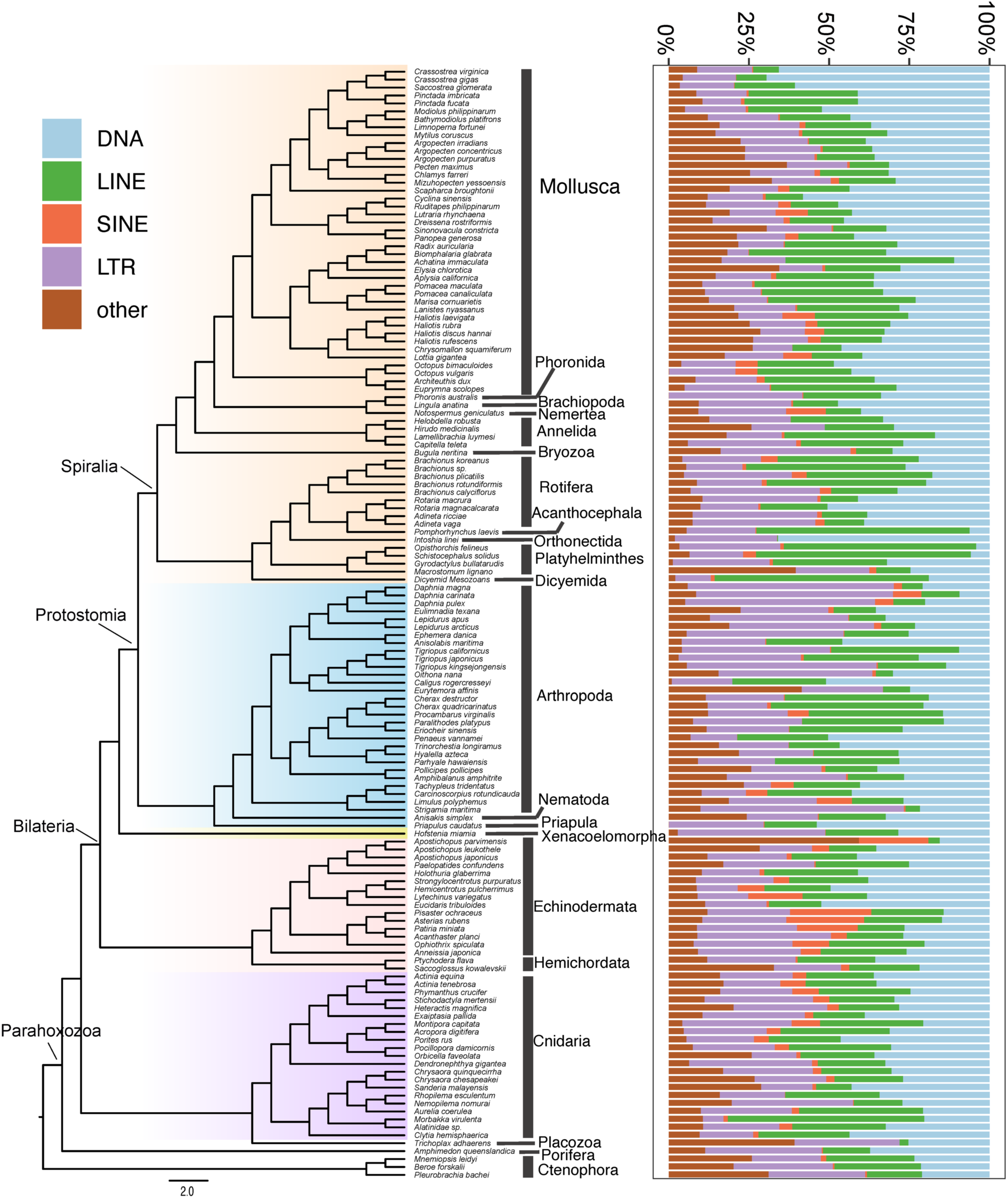
Distribution of repeat subtypes in marine invertebrate genomes.

**Figure S5.**
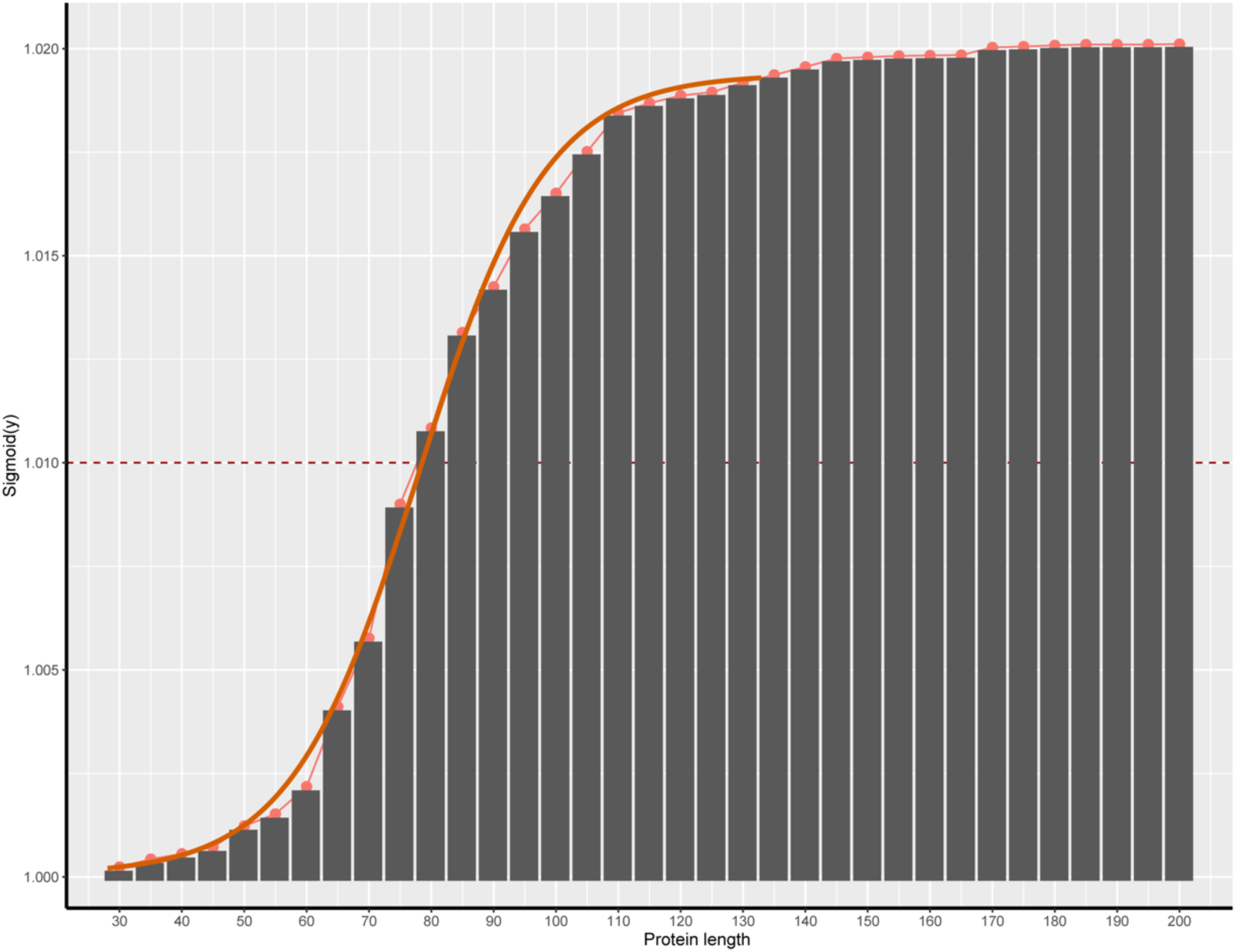
Sigmoid model of homeobox protein length of marine invertebrates.

**Figure S6.**
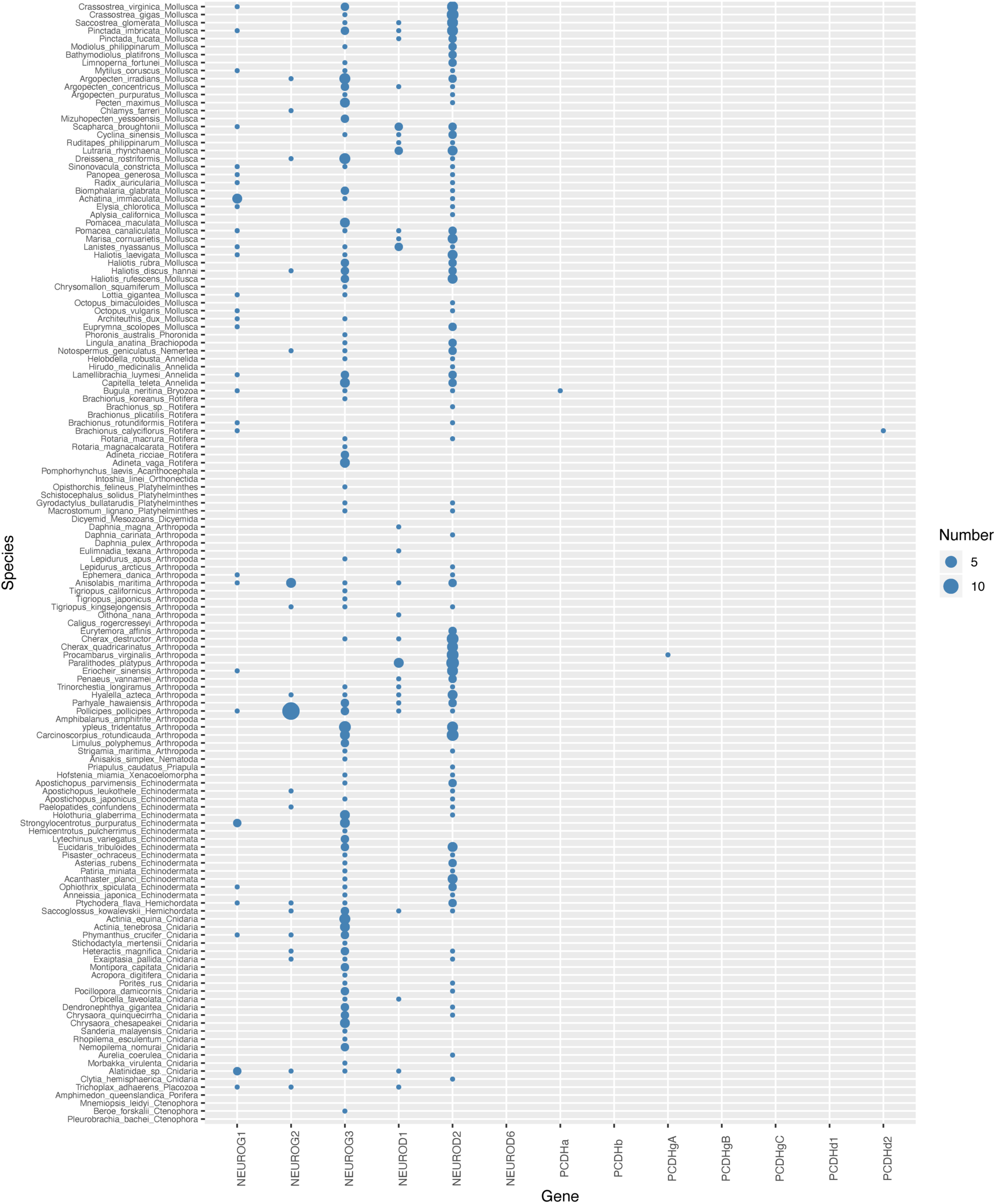
Distribution of identified fragments (less than 80 amino acid) of nervous system related genes.

